# ERFMTDA: Predicting tsRNA–disease associations using an enhanced rotative factorization machine

**DOI:** 10.64898/2026.03.20.713298

**Authors:** Wei Lan, Dong Wang, Wenyi Chen, Xuhua Yan, Qingfeng Chen, Shirui Pan, Yi Pan

## Abstract

**Motivation:** tRNA-derived small RNAs (tsRNAs) have emerged as a novel class of regulatory molecules implicated in the pathogenesis of many human diseases, making them as promising biomarkers and therapeutic targets. However, existing computational methods for tsRNA–disease association prediction often overlook explicit biological attributes and complex feature interactions, limiting their predictive performance.

**Results:** We propose ERFMTDA, an enhanced rotative factorization machine framework for predicting potential tsRNA–disease associations. ERFMTDA explicitly models complex interactions among heterogeneous biological features while integrating latent structural representations derived from the global association matrix. In addition, a biologically informed negative sampling strategy based on motif-level sequence similarity is introduced to improve the reliability of negative samples. Extensive experiments demonstrate that ERFMTDA consistently outperforms eleven state-of-the-art methods. Case studies on diabetic retinopathy and hepatocellular carcinoma further confirm its ability to prioritize biologically meaningful tsRNA–disease associations.

**Availability and implementation:** The source codes and datasets of ERFMTDA are available at https://github.com/lanbiolab/ERFMTDA.

## 1. Introduction

tRNA-derived small RNAs (tsRNAs) have emerged as a distinct and functionally versatile class of regulatory molecules, generated through precise cleavage of mature or precursor tRNAs under stress or pathological conditions (Chu et al., 2022). In addition, a variety of studies have shown that tsRNAs are associated with multiple human diseases (Huang et al., 2020; Xia et al., 2023). For example, tRF-33 has been identified as a tumor suppressor in gastric cancer. Its dysregulation is closely associated with cancer progression, highlighting its potential as a diagnostic and prognostic biomarker (Zhang et al., 2024). These findings indicate that tsRNAs are emerging not only as novel regulators of disease processes but also as promising non-invasive diagnostic biomarkers and therapeutic targets.

Despite the growing recognition of their clinical significance, the functional characterization of tsRNAs remains in its infancy compared to other Non-coding RNAs (ncRNAs). Identifying tsRNA-disease associations through biological experiments is inevitably time-consuming and labor-intensive. Therefore, computational method has become an indispensable strategy for identifying potential disease-related tsRNAs (Lan et al., 2025b, 2024b). Over the past decade, a variety of computational frameworks have been developed for predicting disease related ncRNAs, including microRNAs (miRNAs), long non-coding RNAs (lncRNAs), and circular RNAs (circRNAs) (Lei et al., 2021; Lan et al., 2023). For instance, Yu et al. (2022) proposed a meta-path based method to learns node representations by aggregating information from multiple predefined semantic paths in a miRNA-disease-gene heterogeneous network. Tang et al. (2021) developed a Multi-view Multichannel Attention Graph Convolutional Network (MMGCN) to predict potential miRNA-disease associations. Ding et al. (2021) designed VGAE-MDA for miRNA-disease association prediction by using dual variational graph auto-encoders on miRNA-based and disease-based sub-networks. Li et al. (2024a) devised NAGTLDA for lncRNA-disease association prediction by using a node-adaptive graph Transformer that incorporates network structural encoding. Lan et al. (2022) proposed GANLDA for lncRNA-disease association prediction by using a graph attention network combined with multi-layer perceptron. Zhao et al. (2023) developed MCHNLDA, employing a multi-view contrastive learning framework and a heterogeneous graph attention network with LSTM for lncRNA-disease association prediction. Li et al. (2024b) introduced Bi-SGTAR, which operates on known circRNA-disease pairs and employs a bi-view sparse gating mechanism to assess the reliability and truthfulness of predicted associations. Lan et al. (2024a) developed LGCDA to predict circRNA-disease associations by fusing local structural features and global similarity features. Yuan et al. (2023) proposed iCircDA-NEAE, a method that leverages multiple similarity measures and employs accelerated attribute network embedding with a dynamic convolutional autoencoder to predict circRNA-disease associations. Lan et al. (2025a) proposed CLTDA, a contrastive learning–based method that integrates adaptive singular value decomposition and graph convolutional networks for tsRNA–disease association prediction.

The above methods have achieved promising performance in predicting ncRNA–disease associations. But most of them rely heavily on graph structures and similarity-based information, while ignoring explicit biological features and complex feature interactions, thereby limiting their generalization ability for sparse known associations. This limitation is particularly pronounced for tsRNAs, whose functional roles are closely shaped by their biogenesis and sequence characteristics (Muthukumar et al., 2024).

In order to address these limitations, we develop a tsRNA–disease association prediction framework (ERFMTDA) based on Rotative Factorization Machines (RFM) (Tian et al., 2024). Specifically, ERFMTDA integrates different sources of information by combining intrinsic biological attributes with global structural representations. The biological attributes of tsRNAs and diseases are first encoded and embedded to preserve domain-specific semantic information. Meanwhile, latent global structural features are extracted from the tsRNA–disease association matrix, allowing the model to capture global association patterns. These heterogeneous features are then fed into a feature interaction learning module to capture high-order dependencies among feature fields, and the resulting interaction representations are further enhanced through a modulus amplification mechanism. In addition, we introduce a motif similarity–constrained negative sampling strategy to improve the reliability of negative samples. Finally, the learned tsRNA–disease pair representation is projected into a scalar score to estimate the probability of their association. To evaluate the effectiveness of ERFMTDA, we conducted extensive experiments including 5-fold and 10-fold cross-validation as well as de novo testing. The results consistently demonstrate that ERFMTDA outperforms other existing methods in predicting tsRNA–disease associations. In addition, two case studies on diabetic retinopathy and hepatocellular carcinoma further confirm that ERFMTDA can effectively identify disease-related tsRNAs in real biological scenarios.

## 2. Materials and methods

### 2.1. Overview

The overall workflow of ERFMTDA is summarized in Fig 1, which consists of three stages: (1) feature extraction, (2) feature interaction learning and tsRNA-disease association prediction, (3) negative sampling based on motif similarity.

**Figure 1.**
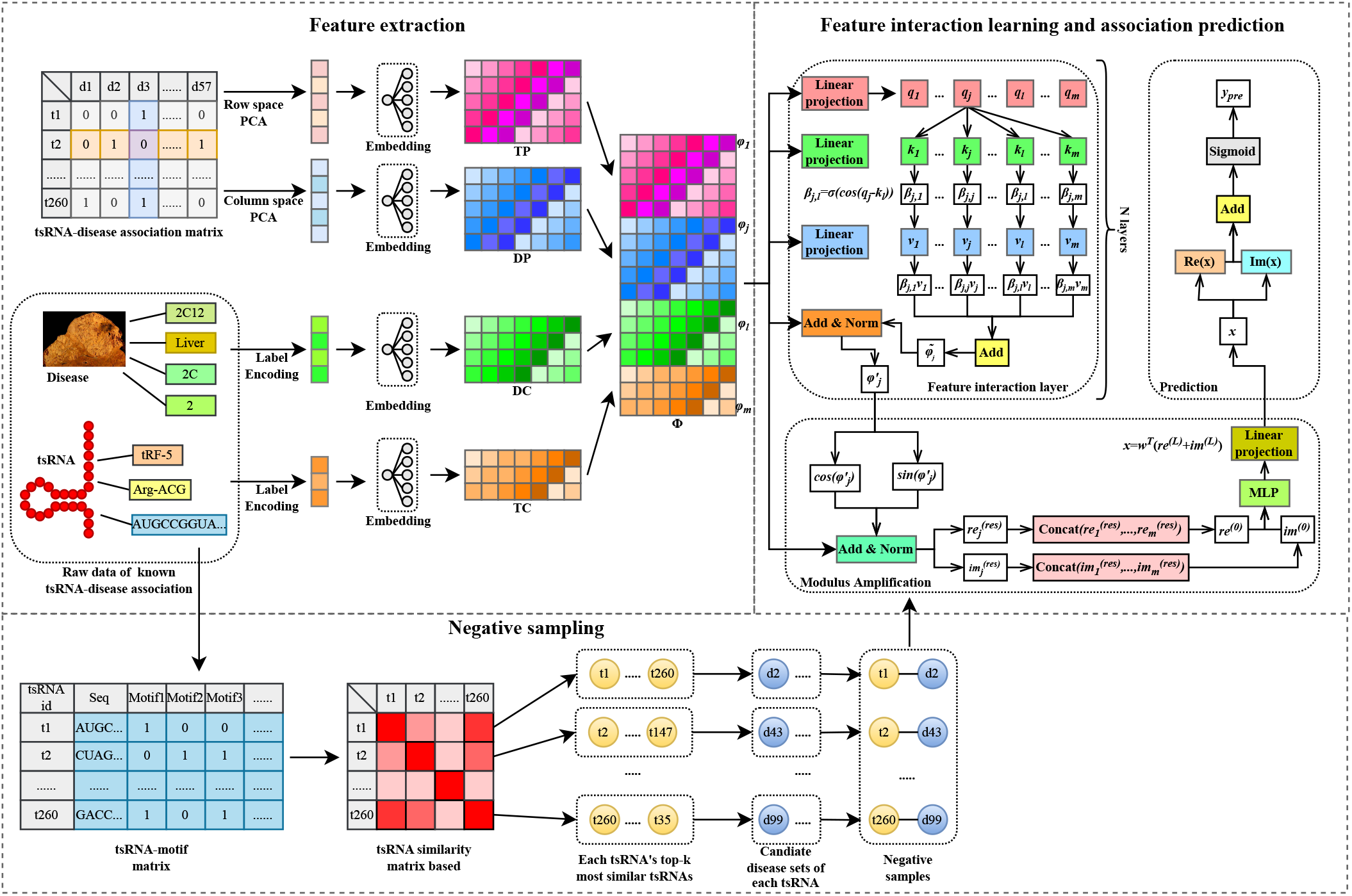
The architecture of ERFMTDA

### 2.2. Data collection

To ensure high data quality and experimental reliability, we constructed a curated tsRNA–disease association dataset through a systematic literature search. Specifically, we searched PubMed using the keywords “tsRNA”, “tRF”, “tRNA-derived fragments”, and “disease”, and manually screened all retrieved publications.

To guarantee the reliability of the collected associations, we applied strict inclusion criteria and retained only tsRNA–disease pairs that were experimentally validated using reverse transcription quantitative PCR (RT-qPCR). After applying this filtering procedure, the final dataset contained 260 tsRNAs, 57 diseases, and 305 experimentally confirmed associations.

### 2.3. Feature extraction

#### 2.3.1. Biological feature encoding and embedding

Each tsRNA contains several intrinsic biological attributes, such as tsRNA type, isotype, and sequence length. These attributes are treated as categorical features and encoded using label encoding, where each category is mapped to a unique integer index. Accordingly, each tsRNA is represented as a discrete feature vector 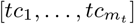, where *m*_*t*_ denotes the number of biological attributes.

Similarly, diseases are characterized by semantic attributes such as ICD codes and affected organs. These attributes are encoded using the same strategy, yielding a discrete representation 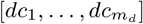, where *m*_*d*_ denotes the number of disease attributes.

To enable effective representation learning, categorical features are mapped into dense vectors via embedding lookup operations. The embedding vectors of the *j*-th tsRNA biological feature and the *k*-th disease semantic feature are defined as:

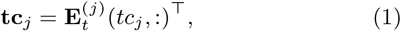

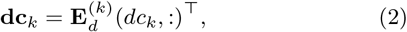

where 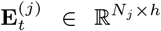 and 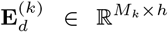 are learnable embedding matrices for the *j*-th tsRNA biological feature and the *k*-th disease semantic feature, respectively. 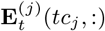 and 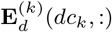 denote the rows of the corresponding embedding matrices indexed by *tc*_*j*_ and *dc*_*k*_. *N*_*j*_ and *M*_*k*_ represent the number of categories for each feature, and *h* denotes the dimensionality of the embedding space.

After embedding, each tsRNA and disease are represented as:

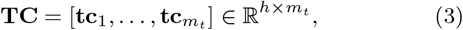

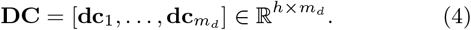

#### 2.3.2. Global structural features extraction

To capture the global interaction structure between tsRNAs and diseases, we construct an association matrix 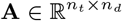, where *n*_*t*_ and *n*_*d*_ denote the numbers of tsRNAs and diseases, respectively:

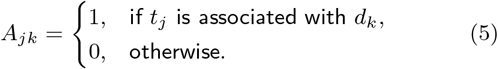

Since the association matrix is typically sparse and high-dimensional, Principal Component Analysis (PCA) is applied to extract compact global structural representations. By projecting the row and column profiles of **A** onto a *u*-dimensional principal subspace, each tsRNA and disease is represented by a *u*-dimensional structural feature vector.

To enable interaction with biological and semantic embeddings, each principal component is further projected into the same *h*-dimensional embedding space. Let *tp*_*l*_ and *dp*_*l*_ denote the *l*-th principal component values of tsRNAs and diseases obtained from PCA, respectively. The corresponding embeddings are defined as:

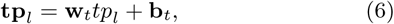

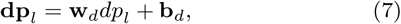

where **w**_*t*_ ∈ ℝ^*h*^ and **w**_*d*_ ∈ ℝ^*h*^ are learnable weight vectors. **b**_*t*_ ∈ ℝ^*h*^ and **b**_*d*_ ∈ ℝ^*h*^ are bias terms. The resulting structural embeddings for tsRNAs and diseases are expressed as:

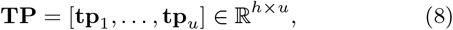

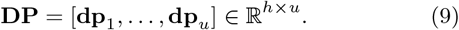

Finally, the biological, semantic, and structural features are concatenated to form a unified representation for each tsRNA–disease pair:

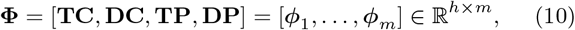

where *m* = *m*_*t*_ + *m*_*d*_ + 2*u* is the total number of feature embeddings.

### 2.4. RFM-based tsRNA-disease association prediction

#### 2.4.1. Feature interaction layer

To capture dependencies among heterogeneous tsRNA and disease features, we model feature interactions using a rotation-based attention mechanism (Tian et al., 2024).

For each feature embedding ***ϕ***_*j*_, query, key, and value representations are obtained through linear projections:

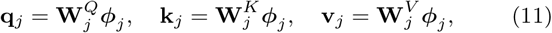

where 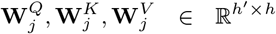 are learnable projection matrices, and *h*^′^ denotes the dimensionality of the projected representations.

To model diverse interaction patterns, a multi-head strategy is employed to split each representation into *D* heads:

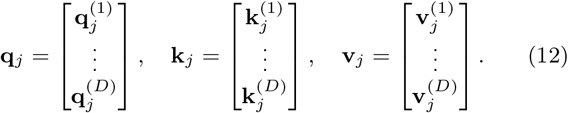

For each head, the relevance between features *j* and *l* is computed based on angular similarity:

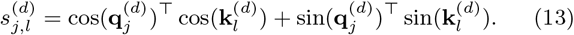

The corresponding attention weight is obtained as:

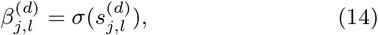

where *σ*(·) denotes the sigmoid function.

Using these weights, the contextual representation of feature *j* is obtained as:

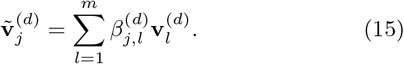

The outputs from all heads are concatenated to form the aggregated representation:

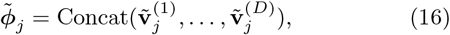

To preserve the original feature information, a residual connection with linear projection is applied:

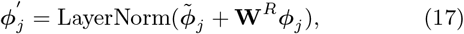

where 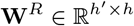 is a shared projection matrix. By stacking multiple such layers, the model progressively refines feature interactions.

#### 2.4.2. Modulus Amplification

After the final interaction layer, each refined feature embedding is mapped to the complex plane as follows:

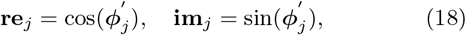

where **re**_*j*_ and **im**_*j*_ denote the real and imaginary components of the *j*-th feature, respectively.

To preserve first-order feature information, residual projections are incorporated:

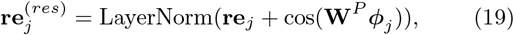

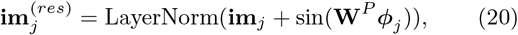

where 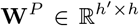 is a learnable projection matrix ensuring dimensional consistency.

The resulting representations encode feature interactions in their angular components, while all features lie on the unit circle with fixed magnitude. However, fixed modulus limits model expressiveness. To address this limitation, the real and imaginary components of all features are concatenated and flattened to form global representations:

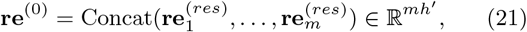

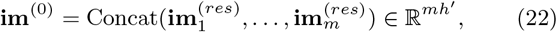

where *mh*^′^ is the dimension of the global representation. Then they are processed by a shared multilayer perceptron to learn adaptive feature-wise amplitudes:

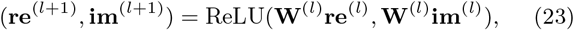

where **W**^(*l*)^ denotes the weight matrix of the *l*-th amplification layer.

After the final amplification layer, the representation is projected to a scalar:

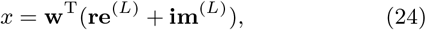

where *L* denotes the number of amplification layers and **w** is a learnable transformation vector.

#### 2.4.3. Prediction and Model Training

The predicted association score between a tsRNA and a disease is obtained as:

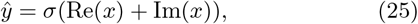

where *σ*(·) is the sigmoid function, Re(*x*) and Im(*x*) are the real part and imaginary part of *x*, respectively.

The model is trained using binary cross-entropy loss with *L*2 regularization:

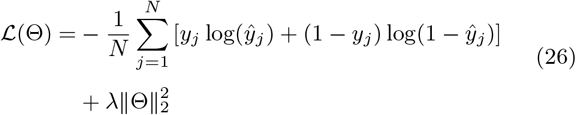

where *y*_*j*_ is the ground truth label. *ŷ*_*j*_ is the predicted value. Θ denotes all trainable parameters, and *λ* is the *L*2-norm penalty.

### 2.5. Negative sampling method

Random negative sampling may introduce noisy instances because some randomly selected tsRNA–disease pairs may correspond to undiscovered true associations. To alleviate this issue, we propose a motif similarity–based negative sampling strategy.

For each tsRNA sequence, all motifs with length *L* are extracted. Motifs appearing at least *k*^′^ times are retained to construct a motif vocabulary *V*_*m*_. Each tsRNA *s*_*j*_ is represented as a binary motif vector:

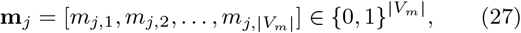

where *m*_*j,k*_ = 1 if motif *v*_*k*_ occurs in *s*_*j*_, and 0 otherwise. Stacking all vectors yields the motif matrix

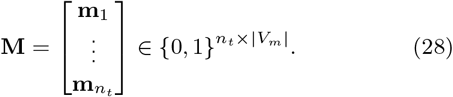

Based on **M**, motif similarity between tsRNAs is computed using cosine similarity:

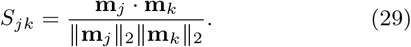

For each tsRNA *t*_*j*_, the top-*k* most similar tsRNAs are collected into the set 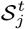. Diseases associated with any tsRNA in 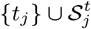 are excluded to form a forbidden disease set:

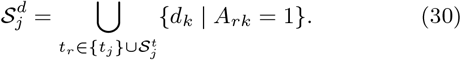

The candidate disease set for *t*_*j*_ is then expressed as:

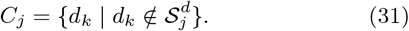

A disease *d*_*k*_ is randomly selected from *C*_*j*_ to construct a negative pair (*t*_*j*_, *d*_*k*_). The procedure is repeated until the number of negative samples equals the number of positive samples.

## 3. Results

### 3.1. Evaluation metrics

To evaluate the predictive performance of ERFMTDA, we performed 5-fold and 10-fold cross-validation, as well as de novo testing.

Six widely used evaluation metrics were adopted, including the area under the receiver operating characteristic curve (AUC), the area under the precision–recall curve (AUPR), accuracy (Acc), precision (Pre), recall (Rec), and F1-score. The definitions of these metrics are provided in the Supplementary Material.

### 3.2. Parameter settings

The key hyperparameters are set as follows: the dimension of feature embeddings and the hidden units in the attention layers are both set to 32. To mitigate overfitting, a dropout rate of 0.1 is applied to both the attention mechanism and the amplification network. The model is optimized using the Adam (Adam et al., 2014) optimizer with a learning rate of 1 × 10^−3^. The weight decay coefficient of *L*2 regularization is set to 1 × 10^−5^. The batch size is set to 32, and the training process is conducted for 200 epochs.

### 3.3. Five-fold cross validation

To evaluate the performance of ERFMTDA, we compare it with eleven representative ncRNA–disease association prediction methods, including IBNPKATZ (Zhao et al., 2019), RWR (Vural et al., 2019), iCircDA-MF (Wei and Liu, 2020), LLCDC (Ge et al., 2020), DWNN-RLS (Yan et al., 2018), RWRKNN (Lei and Bian, 2020), GMNN2CD (Niu et al., 2022), CD-LNLP (Zhang et al., 2019), KATZHCDA (Fan et al., 2018), RNMFLP (Peng et al., 2022), and DMFCDA (Lu et al., 2020). These methods represent several mainstream strategies for ncRNA–disease association prediction, including path-based, matrix factorization–based, kernel-based, and deep learning approaches.

To ensure fair comparison, ERFMTDA and all baseline models are evaluated on the same tsRNA–disease dataset using identical five-fold cross-validation settings. As shown in Fig. 2, ERFMTDA achieves the best overall performance among all competing methods, with an AUC of 0.9004 and an AUPR of 0.9128.

**Figure 2.**
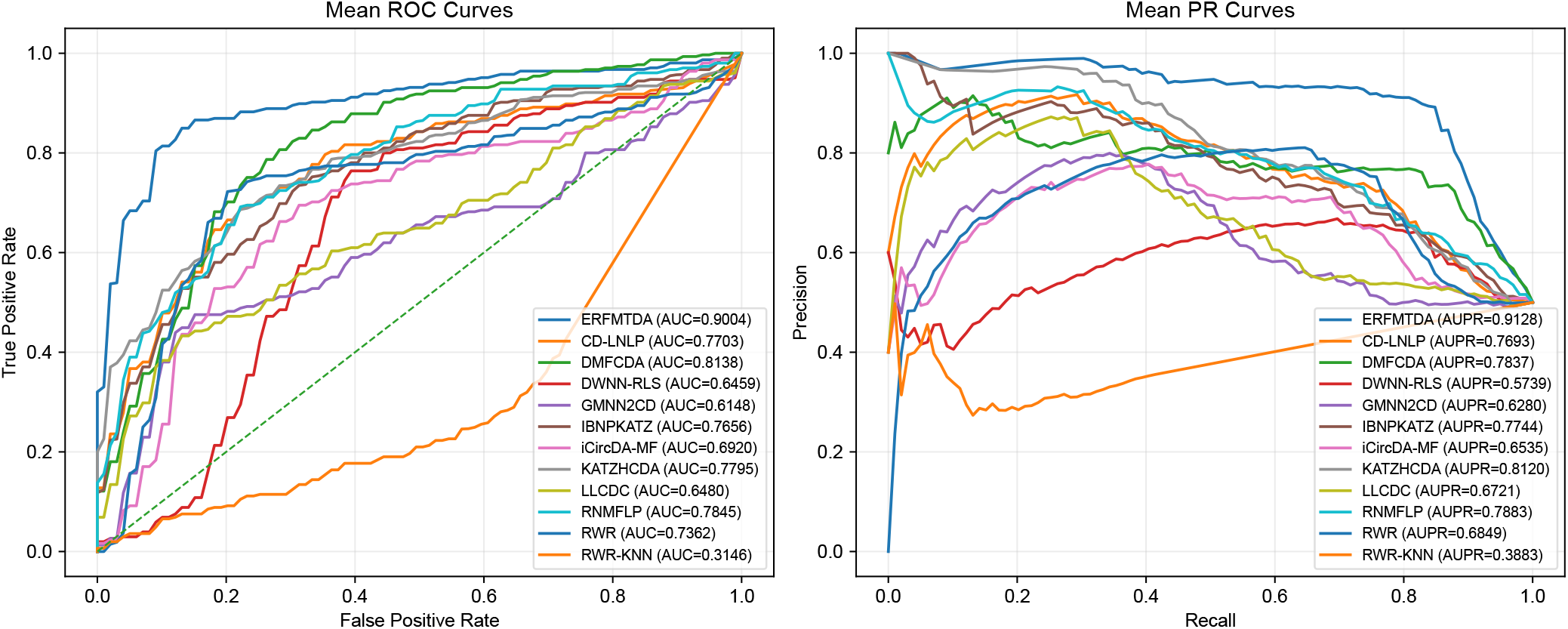
ROC curves and PR curves of ERFMTDA and eleven baseline methods under 5-fold cross-validation.

Compared with the strongest baseline method DMFCDA, ERFMTDA improves AUC and AUPR by 10.6% and 16.5%, respectively. In particular, ERFMTDA shows notable advantages in AUPR, indicating its superior ability to identify tsRNA–disease associations under sparse data conditions. Additional evaluation metrics, including Accuracy, Precision, Recall, and F1-score, are summarized in Table 1, where ERFMTDA consistently achieves the best performance.

**Table 1.**
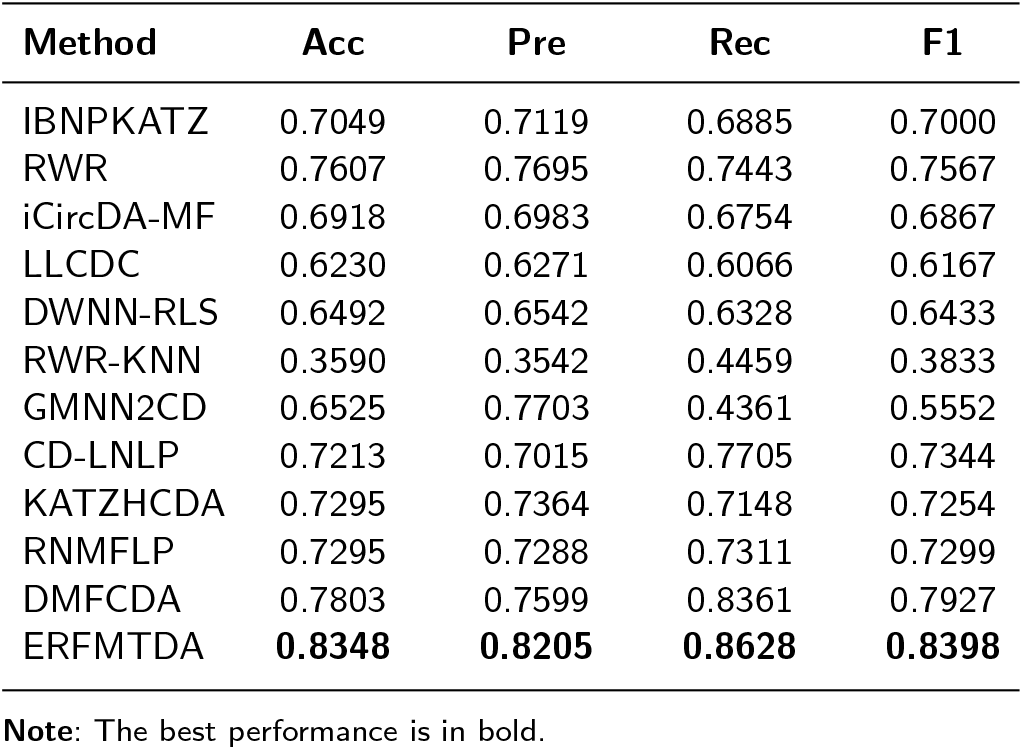
Performance comparison of ERFMTDA with eleven baseline methods under 5-fold cross-validation.

### 3.4. Ten-fold cross validation

To further evaluate model robustness, we additionally conduct ten-fold cross-validation using the same dataset and experimental settings. As shown in Fig. 3, ERFMTDA again achieves the best performance among all competing methods, with an AUC of 0.9009 and an AUPR of 0.9148. The performance trends are consistent with those observed in five-fold cross-validation, and ERFMTDA maintains superior results across all evaluation metrics, as summarized in Table s1 (in the Supplementary Material). These results indicate that the predictive performance of ERFMTDA is stable and not sensitive to different data partition strategies.

**Figure 3.**
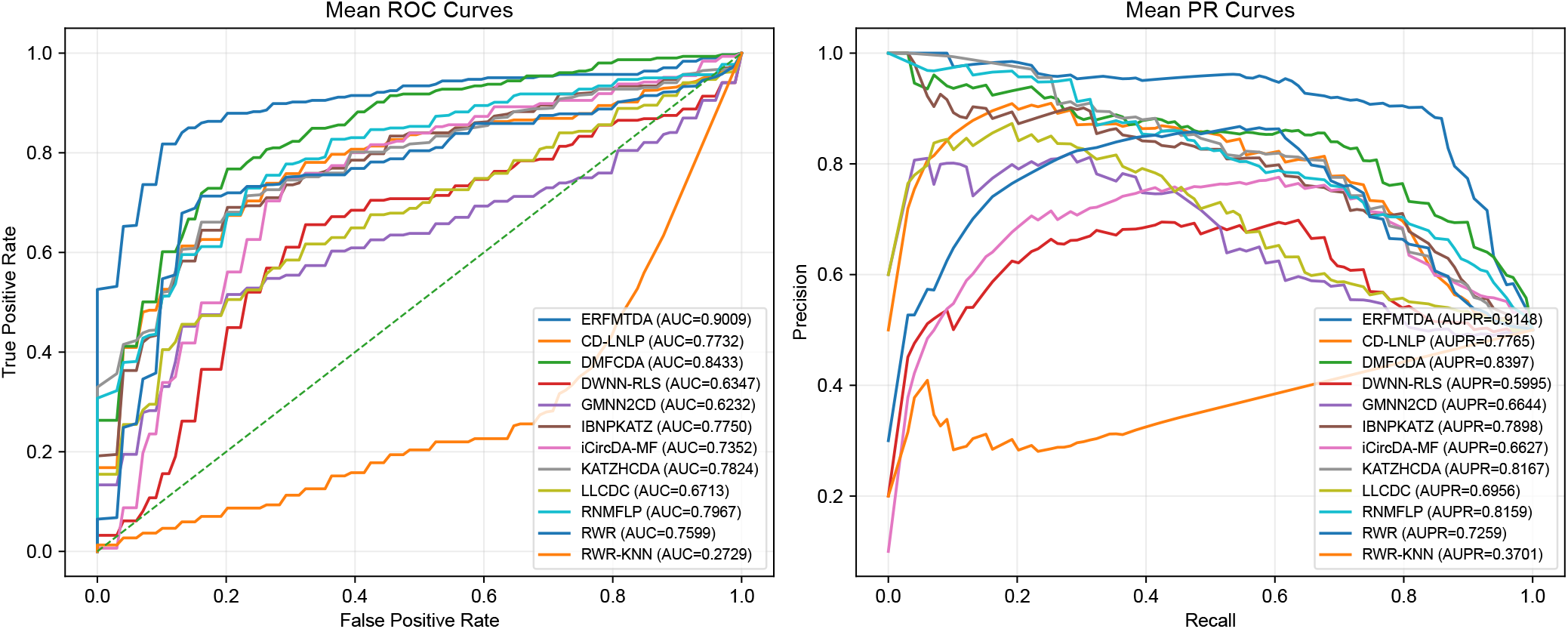
ROC curves and PR curves of ERFMTDA and eleven baseline methods under 10-fold cross-validation.

### 3.5. Component analysis

#### 3.5.1. Ablation study

To evaluate the contributions of different components in ERFMTDA, we construct several ablated variants by removing the global structural features (ERFMTDA/-pca), the motif-similarity-based negative sampling strategy (ERFMTDA/-ns), or both (ERFMTDA/-pca,-ns). As shown in Fig. 4, removing either component leads to noticeable performance degradation. These results indicate that both global structural features and the proposed negative sampling strategy contribute positively to model performance. The full model consistently achieves the best results across all evaluation metrics, demonstrating the effectiveness of integrating these two components.

**Figure 4.**
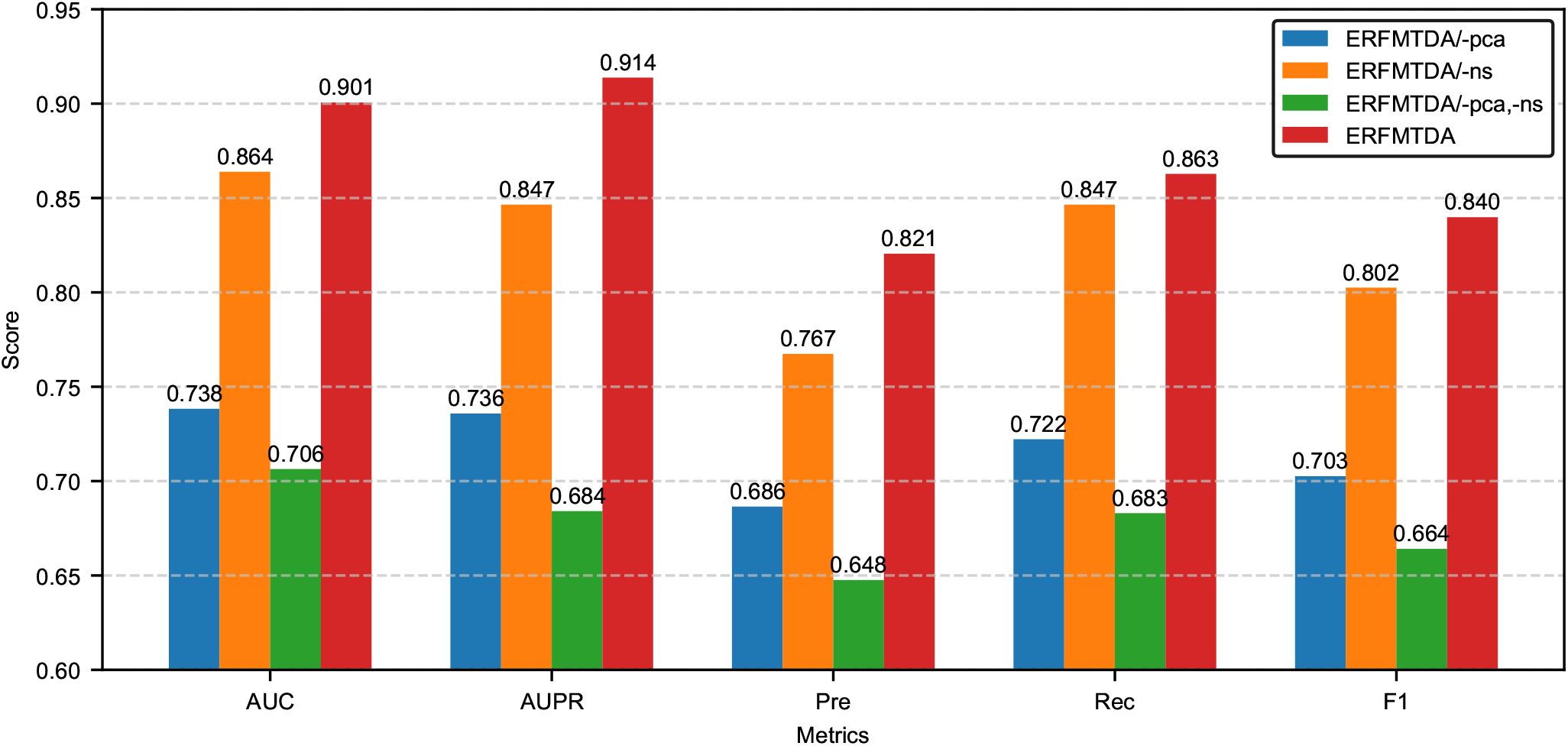
Performance comparison of ERFMTDA and its ablated variants.

#### 3.5.2. Effectiveness analysis of the negative sampling strategy

Random negative sampling may introduce instability due to its inherent randomness, potentially leading to performance variations across different runs. To further evaluate the effectiveness and reliability of the proposed motif-similarity-based negative sampling strategy, we conduct six independent experiments for both ERFMTDA and ERFMTDA/-ns, and report the averaged evaluation metrics in Table s2 (in the Supplementary Material). The full model achieves an average AUC of 0.8858, AUPR of 0.8933, Precision of 0.7986, Recall of 0.8639, and F1-score of 0.8289, all of which are consistently higher than the corresponding averages of ERFMTDA/-ns. These results provide strong evidence that the motif-similarity-based negative sampling strategy effectively enhances model performance.

### 3.6. De novo experiment

To evaluate the ability of ERFMTDA in predicting associations for previously unseen diseases, a de novo experiment was performed. For each disease, all its known tsRNA associations were removed from the training set, and the model was trained on the remaining data to predict associations between the held-out disease and all tsRNAs. This procedure was repeated for every disease.

As shown in Fig. 5, ERFMTDA achieves the best performance among all methods, with an AUC of 0.8116. The detailed evaluation metrics are reported in Table s3 (in the Supplementary Material). These results indicate that ERFMTDA maintains strong predictive capability when encountering unseen diseases, demonstrating its robustness and generalization ability in de novo prediction scenarios.

**Figure 5.**
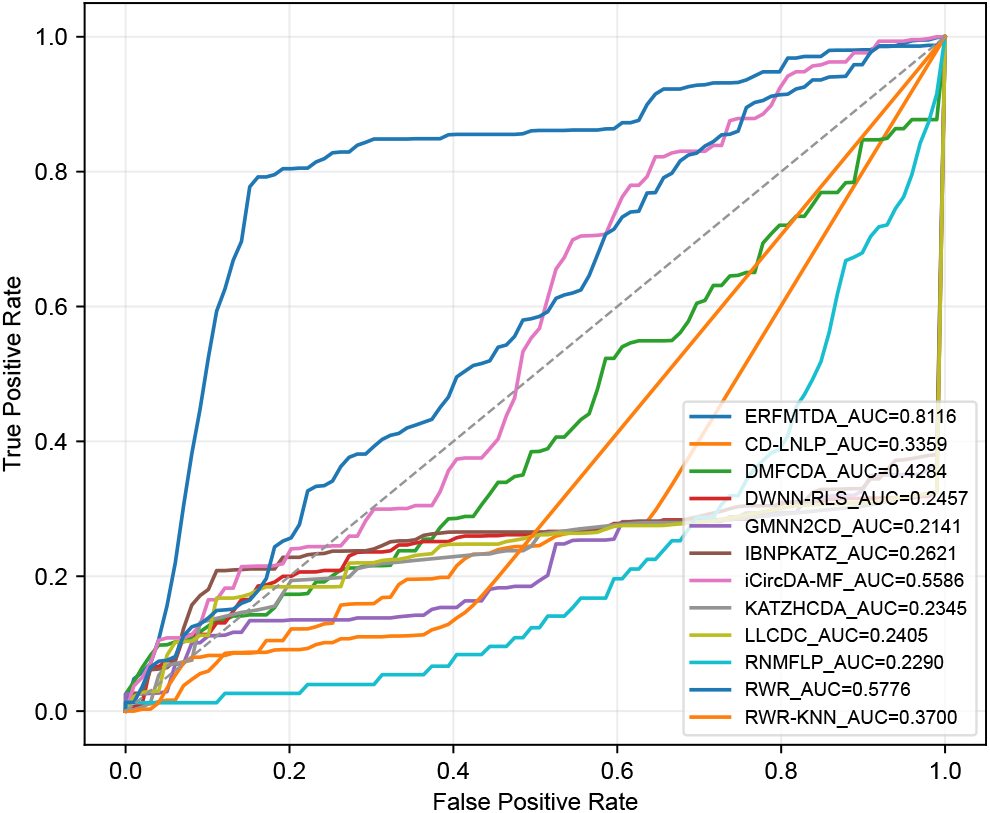
ROC curves of ERFMTDA and other baseline methods in the denovo experiment.

### 3.7. Hyperparameter sensitivity analysis

To investigate the joint effects of the number of principal components *u* in the association matrix and the top-*k* parameter in the negative sampling module, we conduct a hyperparameter sensitivity analysis using AUC and AUPR as evaluation metrics. The results are shown in Fig. 6.

**Figure 6.**
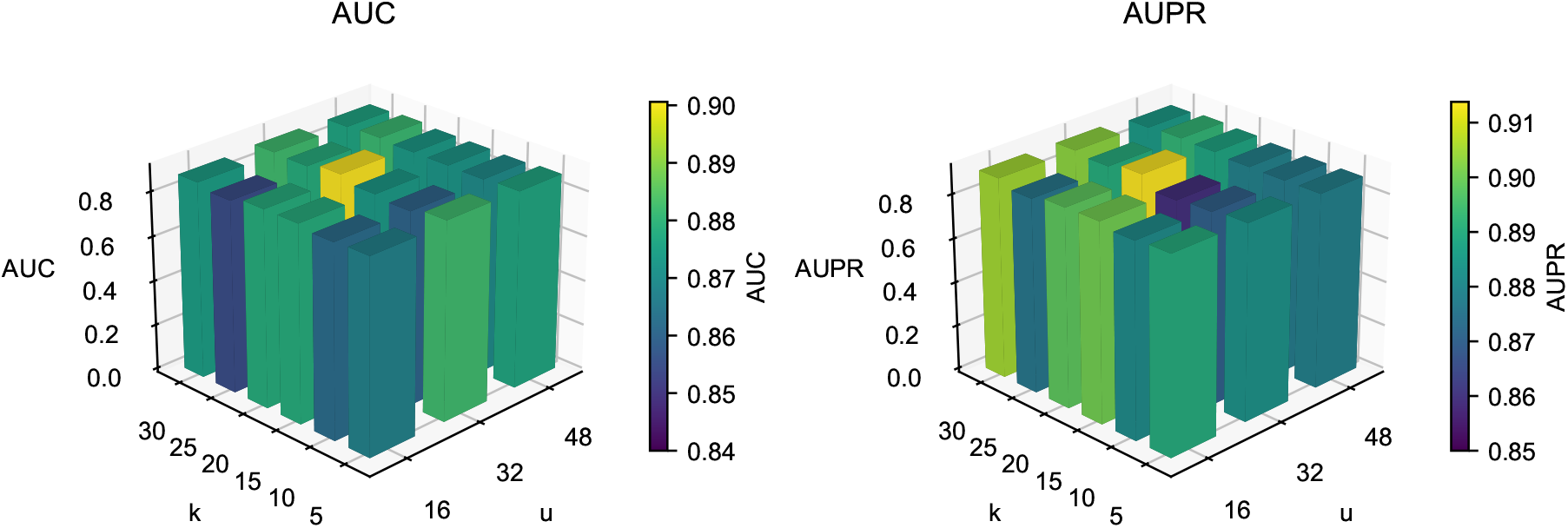
Performance landscapes (AUC and AUPR) under different combinations of *u* and *k*.

ERFMTDA achieves the best performance when *u* = 32 and *k* = 20, yielding the highest AUC (0.9006) and AUPR (0.9138). When *u* is fixed, performance with respect to *k* generally shows an increasing-then-decreasing trend, indicating that moderate *k* values effectively balance the avoidance of false negatives and the diversity of training samples. Similarly, when *k* is fixed, *u* = 32 consistently produces better results than smaller (*u* = 16) or larger (*u* = 48) values, suggesting a suitable trade-off between structural information preservation and noise reduction.

### 3.8. Case study

To further evaluate the practical applicability of ERFMTDA in identifying biologically meaningful tsRNA–disease associations, we conduct two case studies on Diabetic Retinopathy (DR) and Hepatocellular Carcinoma (HCC), which represent two distinct disease contexts.

#### 3.8.1. Diabetic Retinopathy

All tsRNAs are ranked according to their predicted association scores with DR using the trained ERFMTDA model. The top 10 candidate tsRNAs are listed in Table 2. Among them, several tsRNAs have been previously reported to be associated with DR. For example, 5^′^tiRNA-His-GTG has been reported to regulate Müller glial activity and retinal neurovascular dysfunction under diabetic stress (Yao et al., 2025). tiRNA-Val is significantly upregulated in DR retinal tissues and promotes endothelial proliferation via the Sirt1/Hif-1*α* pathway (Xu et al., 2022).

**Table 2.** Top 10 predicted tsRNAs that associated with Diabetic Retinopathy.

| Rank | tsRNA | Evidence |
| --- | --- | --- |
| 1 | 5'tiRNA-His-GTG | PMID: 39513253 |
| 2 | tiRNA-Val | PMID: 35346383 |
| 3 | 5tiRNA-35-PheGAA-8 | PMID: 39167377 |
| 4 | tRF-Glu-CTC | unconfirmed |
| 5 | tiRNA-1:33-Gly-GCC-1 | unconfirmed |
| 6 | tRNA-ValAAC-5 | unconfirmed |
| 7 | tRNA-GlyTCC-5 | unconfirmed |
| 8 | tiRNA-Val-CAC-001 | unconfirmed |
| 9 | tRF- 1020 | PMID: 36128646 |
| 10 | tRF-30 | PMID: 39001618 |

Meanwhile, several predicted tsRNAs, including tRF-Glu-CTC, tiRNA-1:33-Gly-GCC-1, tRNA-ValAAC-5, tRNA-GlyTCC-5, and tiRNA-Val-CAC-001, have not yet been experimentally reported in DR, suggesting that they may represent potential novel biomarkers or therapeutic targets for future investigation.

#### 3.8.2. Hepatocellular Carcinoma

To further examine the generalization ability of ERFMTDA across different disease types, we perform a case study on HCC. The top 10 predicted tsRNAs are shown in Table 3. Among them, several have been reported to play important roles in HCC. For instance, tiRNA-Gly-GCC-002 has been identified as a prognostic factor associated with HCC progression (Wu et al., 2024). Gly-tRF promotes HCC cell migration through the AKT signaling pathway (Zhou et al., 2021).

**Table 3.** Top 10 predicted tsRNAs that associated with Hepatocellular Carcinoma.

| Rank | tsRNA | Evidence |
| --- | --- | --- |
| 1 | tiRNA-Gly-GCC-002 | PMID: 39430835 |
| 2 | CU1276 | unconfirmed |
| 3 | Gly-tRF | PMID: 34537070 |
| 4 | tRF-39 | PMID: 38455567 |
| 5 | 5'-HisGUG | unconfirmed |
| 6 | 5'-tiRNA-Gln | PMID: 36973570 |
| 7 | tRNA-GluCTC-5 | unconfirmed |
| 8 | LeuCAG3'tsRNA | PMID: 29186115 |
| 9 | tRF3b-PheGAA-6 | unconfirmed |
| 10 | tRF3-28-PheGAA-1 | unconfirmed |

In addition, several other predicted tsRNAs, including CU1276, 5^′^-HisGUG, tRNA-GluCTC-5, tRF3b-PheGAA-6, and tRF3-28-PheGAA-1, remain experimentally unconfirmed, indicating that ERFMTDA may provide valuable candidates for future biological validation.

Overall, the case studies demonstrate that ERFMTDA can successfully recover known disease-associated tsRNAs while also identifying novel high-confidence candidates across different disease contexts, highlighting its potential utility for discovering biologically meaningful tsRNA–disease associations.

## 4. Conclusion

In this study, we propose ERFMTDA, a computational framework for predicting tsRNA–disease associations based on Rotative Factorization Machines. By integrating intrinsic biological attributes with latent global structural representations derived from the association matrix, ERFMTDA effectively captures complex feature interactions within a unified embedding space. In addition, a motif-based sequence similarity–guided negative sampling strategy was introduced to improve the reliability of negative samples. Extensive evaluations demonstrate that ERFMTDA consistently outperforms existing approaches. Case studies on diabetic retinopathy and hepatocellular carcinoma further confirm its ability to identify biologically meaningful disease-related tsRNAs.

Despite these encouraging results, several limitations remain. The relatively small number of experimentally validated tsRNA–disease associations restricts the diversity of training data, and the current framework does not explicitly model tsRNA secondary structures. Future work will focus on incorporating additional high-quality association data and integrating structural features to further improve the robustness and biological interpretability of the model.

