## Supplemental Table1, Table2, Table3 for "ERFMTDA: Predicting tsRNA–disease associations using an enhanced rotative factorization machine"

### Part A: Evaluation Metrics

Let TP, TN, FP, and FN denote the numbers of true positives, true negatives, false positives, and false negatives, respectively. The evaluation metrics used in this study are defined as follows:

$$\mathrm{Acc} = \frac{\mathrm{TP} + \mathrm{TN}}{\mathrm{TP} + \mathrm{TN} + \mathrm{FP} + \mathrm{FN}} \tag{32}$$

$$\mathrm{Pre} = \frac{\mathrm{TP}}{\mathrm{TP} + \mathrm{FP}} \tag{33}$$

$$\mathrm{Rec} = \frac{\mathrm{TP}}{\mathrm{TP} + \mathrm{FN}} \tag{34}$$

$$\mathrm{F1} = \frac{2 \times \mathrm{Pre} \times \mathrm{Rec}}{\mathrm{Pre} + \mathrm{Rec}} \tag{35}$$

$$\mathrm{TPR} = \frac{\mathrm{TP}}{\mathrm{TP} + \mathrm{FN}} \tag{36}$$

$$\mathrm{FPR} = \frac{\mathrm{FP}}{\mathrm{FP} + \mathrm{TN}} \tag{37}$$

The ROC curve is constructed by plotting the true positive rate (TPR) against the false positive rate (FPR) under varying classification thresholds, and its area (AUC) reflects the model's overall discriminative ability. The precision–recall (PR) curve plots Precision versus Recall across thresholds, and its area (AUPR) characterizes the balance between these two quantities. Together, these metrics provide a multidimensional evaluation of ERFMTDA's predictive performance.

### Part B: Additional Experimental Results

Table S1: Performance comparison of ERFMTDA with eleven baseline methods under 10-fold cross-validation.

| Method | Acc | Pre | Rec | F1 |
|---|---|---|---|---|
| IBNPKATZ | 0.7224 | 0.7330 | 0.7060 | 0.7178 |
| RWR | 0.7447 | 0.7519 | 0.7299 | 0.7407 |
| iCircDA-MF | 0.7256 | 0.7333 | 0.7092 | 0.7211 |
| LLCDC | 0.6334 | 0.6380 | 0.6170 | 0.6273 |
| DWNN-RLS | 0.6592 | 0.6647 | 0.6428 | 0.6536 |
| RWR-KNN | 0.2885 | 0.2813 | 0.2721 | 0.2766 |
| GMNN2CD | 0.6599 | 0.7911 | 0.4442 | 0.5609 |
| CD-LNLP | 0.7185 | 0.6989 | 0.7679 | 0.7317 |
| KATZHCDA | 0.7417 | 0.7451 | 0.7382 | 0.7411 |
| RNMFLP | 0.7398 | 0.7388 | 0.7414 | 0.7401 |
| DMFCDA | 0.7738 | 0.7494 | 0.8294 | 0.7858 |
| ERFMTDA | **0.8495** | **0.8441** | **0.8691** | **0.8545** |

**Note**: Acc, Pre, Rec, and F1 of different methods in the 10-fold cross validation experiment. The best performance is in bold.

Table S2: Average performance of ERFMTDA and ERFMTDA/-ns over six independent experiments.

| **Method** | **Seed** | **AUC** | **AUPR** | **Pre** | **Rec** | **F1** |
|---|---|---|---|---|---|---|
| ERFMTDA/-ns | 7 | 0.8619 | 0.8347 | 0.7872 | 0.8825 | 0.8303 |
| ERFMTDA/-ns | 42 | 0.8622 | 0.8618 | 0.7735 | 0.8464 | 0.8081 |
| ERFMTDA/-ns | 512 | 0.8557 | 0.8539 | 0.7820 | 0.8693 | 0.8229 |
| ERFMTDA/-ns | 1024 | 0.8678 | 0.8727 | 0.7640 | 0.8336 | 0.7957 |
| ERFMTDA/-ns | 1983 | 0.8639 | 0.8465 | 0.7674 | 0.8465 | 0.8025 |
| ERFMTDA/-ns | 2025 | 0.8766 | 0.8910 | 0.8048 | 0.8660 | 0.8335 |
| ERFMTDA/-ns(mean) | / | 0.8647 | 0.8601 | 0.7798 | 0.8574 | 0.8155 |
| ERFMTDA | 7 | 0.9122 | 0.9152 | 0.8258 | 0.8792 | 0.8511 |
| ERFMTDA | 42 | 0.8845 | 0.9023 | 0.8118 | 0.8823 | 0.8453 |
| ERFMTDA | 512 | 0.8731 | 0.8812 | 0.7562 | 0.8594 | 0.8037 |
| ERFMTDA | 1024 | 0.8670 | 0.8547 | 0.7794 | 0.8631 | 0.8177 |
| ERFMTDA | 1983 | 0.9006 | 0.9138 | 0.8205 | 0.8628 | 0.8398 |
| ERFMTDA | 2025 | 0.8776 | 0.8928 | 0.7978 | 0.8366 | 0.8156 |
| ERFMTDA(mean) | / | **0.8858** | **0.8933** | **0.7986** | **0.8639** | **0.8289** |

**Note**: The mean performance of ERFMTDA/-ns is underlined, while the mean performance of ERFMTDA is highlighted in bold.

Table S3: Performance comparison of ERFMTDA and other baseline methods in the de novo experiment.

| **Method** | **AUPR** | **Pre** | **Rec** | **F1** |
|---|---|---|---|---|
| IBNPKATZ | 0.0398 | 0.0075 | 0.2656 | 0.0137 |
| RWR | 0.0548 | 0.0240 | 0.5810 | 0.0433 |
| iCircDA-MF | 0.0478 | 0.0139 | 0.5633 | 0.0262 |
| LLCDC | 0.0388 | 0.0069 | 0.2615 | 0.0127 |
| DWNN-RLS | 0.0303 | 0.0074 | 0.2622 | 0.0135 |
| RWR-KNN | 0.0251 | 0.0142 | 0.6005 | 0.0268 |
| GMNN2CD | 0.0444 | 0.0065 | 0.1955 | 0.0118 |
| CD-LNLP | 0.0265 | 0.0078 | 0.1305 | 0.0130 |
| KATZHCDA | 0.0305 | 0.0072 | 0.2665 | 0.0132 |
| RNMFLP | 0.0437 | 0.0172 | 0.1409 | 0.0296 |
| DMFCDA | 0.0673 | 0.0168 | 0.1792 | 0.0186 |
| ERFMTDA | **0.1149** | **0.0358** | **0.8596** | **0.0646** |

**Note**: AUPR, Pre, Rec, and F1 of different methods in the de novo experiment. The best performance is in bold.
